# Partial protection from fluctuating selection leads to evolution toward wider population size fluctuation and a novel mechanism of balancing selection

**DOI:** 10.1101/2022.07.08.499270

**Authors:** Yuseob Kim

## Abstract

Classical theory predicted that natural selection favors a variant causing smaller fluctuation of population density. It is also known that if population is partially protected from fluctuating selection, as in the case of seed bank, the variance of fitness is further reduced and therefore the reproductive success of population is ensured. This study, exploring a mathematical model for coupled demographic and evolutionary dynamics, finds that such a ‘refuge’ from fluctuating selection even causes positive selection for a variant increasing the amplitude of population size fluctuation under weak or moderate regulation of population density. Under strong density regulation and constant carrying capacity, long-term maintenance of polymorphism known as the storage effect emerges. However, if the carrying capacity is changing either cyclically or randomly and the density regulation is strong, variants whose fitness fluctuates in phase with population size are positively selected to either fixation or oscillation at intermediate frequencies. The latter dynamics, arising when fitness fluctuates as expected under a simple life-history trade-off, is a novel form of balancing selection. These results highlight the importance of allowing in models the joint demographic and population genetic changes, the failure of which prevents the discovery of important and novel eco-evolutionary dynamics.

## Introduction

Natural populations face cyclic or random fluctuations of their abiotic and biotic environment that lead to fluctuation in individuals’ fitness (Grant and Grant 2002; Bell 2010; Thompson 2013). Evolution under fluctuating selection has been a major topic in population genetics. Theoretical models have explored the effect of fluctuating selection on various aspects of evolution such as the level of both neutral and non-neutral genetic variation (Dempster 1955; Haldane and Jayakar 1963; Hedrick 1976; Hedrick 1986; Gillespie 1994; Barton 2000; Huerta-Sanchez *et al*. 2008). Recently, balancing selection under seasonal fluctuation of fitness has received particular attention as a potential explanation for the apparent seasonal oscillation of allele frequencies at thousands of nucleotide sites in temperate *Drosophila melanogaster* populations (Bergland et al. 2014; Wittmann et al. 2017; Machado et al. 2021). One area of theoretical development focuses on the effect of a subset of the population or a particular life cycle stage protected from fluctuating selection in increasing species or allelic diversity (Chesson and Warner 1981; Ellner and Hairston 1994; Gulisija and Kim 2015; Svardal et al. 2015; Bertram and Masel 2019), which is commonly called the storage effect. Simple mathematical models showed that negative frequency-dependent selection emerges in the presence of a refuge from selection, such as a seed bank (Turelli et al. 2001). The storage effect can generate stable long-term polymorphism at multiple loci (Park and Kim 2019), although it requires strong selection (a large magnitude of fitness oscillation; Bertram and Masel (2019)).

Partial protection from fluctuating selection is expected not only with seed banks but also in other conditions commonly encountered by plants and animals. The survival of individuals in diapause, for example in the insect pupal stage, is probably unaffected by a variable environment (Hedrick 1995). In species with overlapping generations, only a portion of a population will be subject to selection on a trait that is expressed at a particular stage of the life-cycle (Chesson and Warner 1981; Ellner and Hairston 1994). In dioecious species, alleles affecting female reproductive traits, for example, are protected from selection if they are carried by males (Reinhold 2000). If an allele conferring genetic robustness or phenotypic plasticity is segregating in a population, it constitutes a genetic background on which new mutants are protected from selection (Gulisija et al. 2016). In all of these cases, a species effectively consists of a subpopulation exposed to selection and another subpopulation, the refuge, protected from selection leading to the storage effect. Individuals (i.e., alleles) move between these subpopulations with a certain probability per unit time.

Temporally variable environments most likely cause a fluctuation not only in the relative fitness among alleles but also their absolute fitness, the per-capita rate of reproduction per unit time. Namely, the evolutionary change of population under fluctuating environment must occur while population size is changing concurrently. If population density regulation, which is defined as the force that pushes the population size toward a carrying capacity set by environment conditions at a given time, is not very strong, population size can also change (e.g. overshoot or undershoot the carrying capacity) as its genetic composition changes. However, theoretical models for the storage effect have considered an infinitely large or constant-sized population, under which the evolutionary dynamics are fully described by the relative fitness and the relative frequencies of alleles. In this study, the storage effect is modeled with absolute fitness, which allows joint changes in demography and allele frequencies. Then, it can be investigated whether and how evolutionary changes interact with fluctuations in population size. First, it remains to be examined whether the maintenance of polymorphism by the storage effect still occurs under variable population sizes due to relaxed density regulation. Second, one may ask whether the nature of population size fluctuation is altered by evolutionary process. The proximate cause of population size fluctuation, observed in a wide range of species in nature, is probably the fluctuation of abiotic and biotic factors in the environment. However, the degree of such demographic fluctuation depends on the phenotypes of individuals that are the product of evolution.

Classical studies in population genetics found that the long-term reproductive success of a population or an allele under fluctuating environment is given by the geometric mean of its fitness over time. The geometric mean decreases as fitness fluctuates in time even if its arithmetic mean remains constant. Therefore, it is predicted that a genotype causing less fluctuation in absolute fitness, for example by allocating resource for survival during a harsh period, is favored under fluctuating environment (Gillespie 1977; Lande et al. 2009). It should be noted that seed bank and other means of partial protection from fluctuating selection were proposed as a means of increasing the geometric mean fitness of a population by reducing the variance of absolute fitness, a mechanism known as the evolutionary bet-hedging (Seger and Brockmann 1987; Philippi and Seger 1989).

Assuming that there are two alleles whose absolute fitness fluctuate differently but yield the same arithmetic or geometric mean fitness outside the refuge, I found that an allele with a wider fluctuation in absolute fitness can be positively selected under weak or moderate regulation of population density. Namely, directional, not negative frequency-dependent, selection occurs and results in the amplification of population size fluctuation. More surprisingly, preexisting oscillation in population size was found to generate positive selection for an allele whose relative fitness oscillates in phase with population size oscillation, leading to its fixation or long-term oscillation at intermediate frequencies. The latter is a novel mechanism of balancing selection that may explain the seasonal oscillation of allele frequencies at multiple loci in Bergland *et al*. (2014).

The main results will be given in two steps. The full model will be first explored using numerical iterations, which reveals various patterns of eco-evolutionary dynamics. Then, theoretical explanations for these results will be sought by mathematical analyses on reduced models.

## Modeling the fitness of a variant under fluctuating selection

Consider a population of haploid individuals reproducing in discrete generations in an environment that cyclically (e.g., seasonally) changes with a period of *λ* generations. In the following, analyses will focus on the fate of a focal allele, *A*_2_, at a locus that arose recently by mutation so that its frequency, *q*, is low initially. The ancestral (wild-type) allele of the locus is *A*_1_ with frequency *p* = 1 - *q*. A trade-off is assumed in the effect of *A*_2_ on fitness over different phases of an environmental cycle. For example, due to the allocation of finite resource over time, if *A*_2_ increases fitness in one season, it inevitably leads to a decline in fitness in the other season. Here, the fitness of an *A*_2_-carrying individual can be defined relative to that of an *A*_1_-carrying individual or as the expected number of offspring per capita (the absolute fitness). In the (implicit) assumption of strictly constant-sized population, one only needs to specify the relative fitness of alleles in the model. However, in order to model the joint dynamics of evolutionary and demographic changes, absolute fitness needs to be specified. Here the model of constant (or infinite) sized population will be first considered.

First, the fitness of *A*_2_ relative to *A*_1_ may be given by *w*_2_ = *e^s_t_^* (*w*_1_ = 1), where

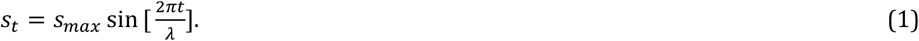

This ensures that the geometric mean fitness of *A*_2_ is equal to that of *A*_1_, because the relative frequencies of *A*_1_ and *A*_2_ after one cycle of fluctuating environment is given by *p’* and *q’*, where

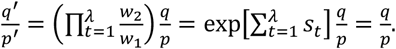

Therefore, allele frequencies are not expected to change in this population. in this case, *A*_1_ and *A*_2_ may be called quasi-neutral to each other. There are other ways to specify the fitness of *A*_1_ and *A*_2_ so that their geometric mean fitness are identical. For example, Gulisija and Kim (2015) assumed *w*_1_ = 1 – *s_t_* and *w*_2_ = 1 + *s_t_* and Bertram and Masel (2019) also used an analogous model where the ratio of two alleles’ fitness is simply reversed upon a seasonal change. This class of fitness assignments may be called geometric mean-preserving fitness-fluctuation, or GMF in short. In the following a mutant allele having this fitness effect will be called a GMF allele.

Second, one may assume *w*_1_ = 1 and *w*_2_ = 1 + *s_t_*, making the arithmetic mean fitness identical. Here, fitness ratio is not reversed in the opposite phase of a cycle (e.g., 1 + *s* < 1/(1 – s)). In this case, it can be easily shown that *q’/p’* < *q/p* after one environmental cycle with *s_max_* > 0. Namely, the derived allele *A*_2_ will be eliminated from the population since its geometric mean fitness is lower than that of *A*_1_. This fitness assignment is called arithmetic mean-preserving fitness-fluctuation or AMF. A mutant allele with this fitness effect will be called an AMF allele.

Which fitness model, GMF or AMF, should be assumed is not a trivial matter as the outcomes of following models critically depend on it. Which one is more realistic and thus should arise more often needs to be addressed as well. Under the strict density regulation in a constant-sized population, *w*_2_/*w*_1_ is approximately the absolute fitness of *A*_2_ when it is rare. Then, AMF means that a fitness gain, i.e. an excess in the number of offspring, in one phase is just enough to compensate the loss in another phase of the environmental cycle. This may happen naturally if a finite resource limits the total number of offspring born in one environmental cycle. In case of GMF, the arithmetic mean of *w*_2_/*w*_1_ over the cycle is greater than 1 (e.g., (*e^s^* + *e^−s^*)/2 > 1 for *s* > 0). Thus, modeling with a GMF allele means that the carrier of a rare derived allele is assumed to produce more offspring, averaged over a cycle, than that of the wild-type allele. One may argue that such a positive effect on fitness may not be achieved by simply changing resource allocation over time. However, there can be many other ways in which fitness fluctuation arises (see more in Discussion). Why the frequency of *A*_2_ does not increase with GMF, despite an increased production of offspring, was well explained by Gillespie (1977) and others who showed that the outcome of fluctuating selection is determined by the geometric mean fitness, which decreases as the variance of fitness fluctuation increases: *A*_2_ at low frequency has higher mean but larger variance in fitness compared to *A*_1_.

## Storage effect under the fluctuation of relative fitness

Next, I will review the theory of the storage effect (Chesson and Warner 1981; Gulisija and Kim 2015; Bertram and Masel 2019). A GMF allele, *A*_2_, is initially assumed in the fluctuating environment: *w*_2_ = *e^s_t_^* and *w*_1_ = 1. The population is now divided into two with fractions 1-*r* and *r*. Only the first subpopulation resides in an environment subject to the fluctuating selection, which I will call the “field”. In the second subpopulation, individuals have identical fitness regardless of alleles, and thus are protected from fluctuating selection. While this “refuge” from selection primarily models a seed bank, it may also correspond to a certain stage of a life cycle in species with overlapping generations, a spatially separated population with relaxed selection, or one sex that is not the target of sex-limited selection. Many of such scenarios are true for diploid species, but the current haploid model should be applicable to diploids with incomplete dominance. It is also assumed that individuals are redistributed into the field and the refuge, with probability 1-*r* and *r*, respectively, during the migration step of each generation. Let *q_t_* be the frequency of *A*_2_ at generation *t*. It is again assumed that *A*_2_ recently arose by mutation and therefore *q_t_* is close to 0. After one cycle of fluctuating selection, frequency *q*_*t*+λ_ is expected to be larger than *q_t_* because

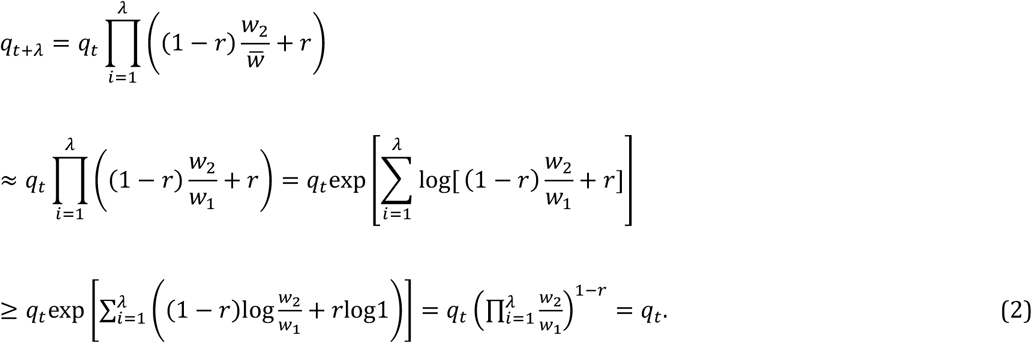

The mean fitness in the field 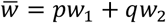 is approximated by *w*_1_ with *q* ≪ *p* and Jensen’s inequality was applied above. No change in allele frequency (*q_t+λ_* = *q_t_*) occurs if fluctuating selection is turned off (*s*_max_ = 0). Therefore, the presence of a refuge confers an advantage to a rare allele under fluctuating selection, thus generating the storage effect.

Furthermore, the above analysis suggests that, to generate the storage effect, the change of environment in the field (fluctuation of *w*_2_ versus *W*_1_) does not have to be periodic as long as the product of 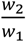 over successive generations converges to 1 with GMF. To confirm this, the iteration of allele frequencies according to the model above was made using *s_t_* = *s_max_*sin[2*πt*\*/Λ], where *t*\* is a random integer between 1 and *λ* drawn each generation. Interestingly, the rare allele increased in frequency slightly faster on average under this mode of random fluctuating selection than under the periodic (“unscrambled”) fluctuation (Fig. 1).

**Figure 1.**
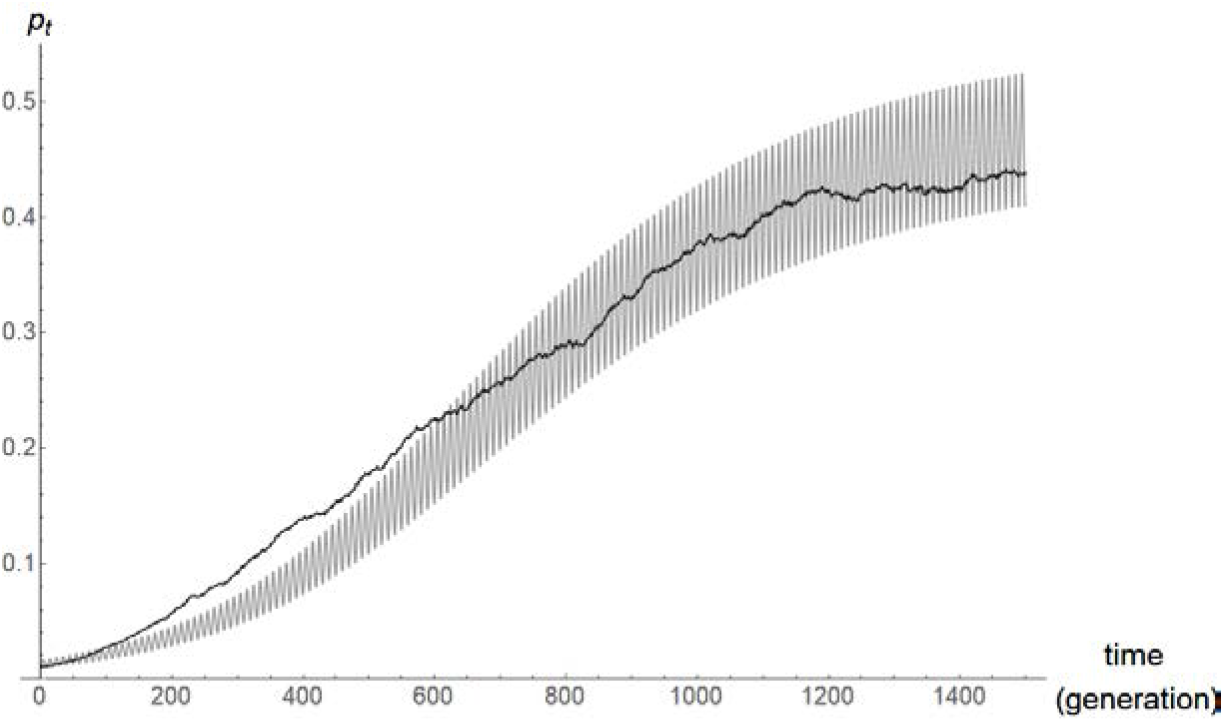
Frequency (*q_t_*) of the *A*_2_, a GMF allele, over time *t* (in generations) under periodic (gray curve; using *s_t_* = *s_max_* sin[2*πt/λ*]) or random (dark curve; using *s_t_* = *s_max_* sin[2*πt^*^/λ*], averaged over 300 replicates) fluctuation of fitness in a population subdivided into the field and the refuge, under the relative fitness-based model. Other parameter: *r* = 0.5, *λ* = 10, *q*_0_ = 0.01, *w*_1_ = 1, *w*_2_ = *e^st^*, and *s_max_* = 0.3.

The storage effect is intriguing because a rare allele is positively selected simply by moving between the field and refuge randomly, even though it is not favored in either subpopulation. It is even possible that a slightly deleterious allele (i.e., having a smaller geometric mean fitness than the other allele in the field) is positively selected when it is rare (Gulisija and Kim 2015). A plain explanation for this phenomenon was that the loss of an allele from the field during an unfavorable period is buffered by its presence in the refuge; then, during the next period when the same allele is now favored in the field, the refuge supplies this allele to the field. However, the buffering effect of a refuge should apply to the other (common) allele as well.

A better explanation may be given by the theory of evolutionary bet-hedging (Seger and Brockmann 1987; Philippi and Seger 1989), according to which a refuge such as a seedbank reduces the variance of reproductive success (of the total population) in a variable environment, thus increasing the geometric mean fitness. This effect of refuge should be larger for the carriers of the rare allele; as mentioned earlier, with GMF, the arithmetic mean of *A*_2_’s fitness is higher than that of *A*_1_ but its fluctuation larger than that of *A*_1_ prevents it from attaining higher geometric mean fitness than *A*_1_. Eq. (2) suggests that, if the variance of fitness in the field is *V*, that of the total population is reduced to (1 – *r*)^2^*V* in the presence of refuge. Although the mean fitness of *A*_2_ also decreases due to refuge (e.g., fitness advantage over *A*_1_ decreases by a factor *r*), this effect must be smaller compared to the decrease of variance; the net effect is to elevate the geometric mean fitness of *A*_2_ over 1. On the other hand, refuge has little effect on the geometric mean fitness of *A*_1_, the major allele, because its absolute fitness fluctuates negligibly. This explains how negative frequency-dependent selection, the storage effect, arises due to partial protection from selection if GMF is assumed.

The storage effect does not arise for an AMF allele (*w*_1_ = 1 and *w*_2_ = 1 + *s_t_*). Again, a refuge reduces the variance of *A*_2_’s fitness and thus increases its geometric mean fitness when *A*_1_ is rare. However, the geometric mean fitness of *A*_2_ relative to *A*_1_ will approach only up to its arithmetic mean fitness relative to *A*_1_, which cannot exceed 1 under the assumption of AMF. Therefore, the proposal of storage effect as a mechanism of balancing selection in the previous studies critically depended on assuming a GMF, not AMF, allele in modeling fluctuating selection.

## Cyclic fluctuation of absolute fitness: Constant carrying capacity

The above model based on relative fitness and relative allele frequencies does not assume any particular population dynamics. However, a fluctuating environment is likely to generate fluctuation in population size as well if population density is not strictly regulated. Furthermore, the dynamics of allele frequency changes may depend on whether population size varies in time or not. I therefore expanded the above model to allow the sizes of subpopulations to change according to the absolute fitness of individuals under varying degrees of population density regulation.

Let *n*_1*f*_ and *n*_2*f*_ (*n*_1*r*_ and *n*_2*r*_) be the numbers of *A*_1_ and *A*_2_ individuals in the field (in the refuge). In the absence of density regulation, *n*_1*f*_ and *n*_2*f*_ are assumed to increase to *W*_1*f*_*n*_1*f*_ and *W*_2*f*_*n*_2*f*_ over a generation in the field. Namely, *W*_1*f*_ and *W*_2*f*_ are the absolute fitness of *A*_1_ and *A*_2_, which may change over time. Absolute fitness for both alleles in the refuge is given by *W_r_*. For density regulation, I define the carrying capacities of the field and refuge to be *K_f_* and *K_r_*, which may also change over time. In addition, it is assumed that, after steps of growth and density regulation, individuals in the field migrate to the refuge with probability *m_fr_* and those in the refuge migrate to the field with probability *m_rf_*. Then, the numbers of *A*_1_ and *A*_2_ individuals in the field and refuge in the next generation become

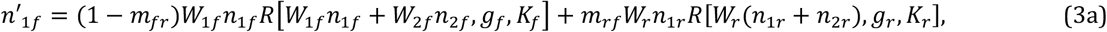

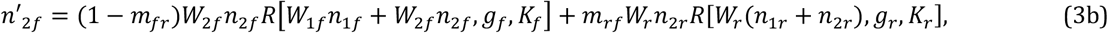

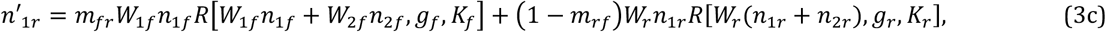

and

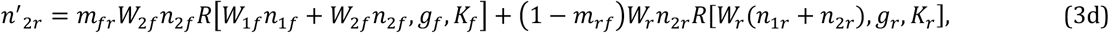

where

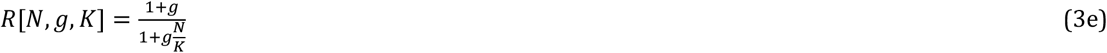

determines the strength of density regulation. If individuals carry only neutral alleles and the population size *N* is much smaller than its carrying capacity *K*, the population will grow at rate *g* (≥ 0). Therefore, a large value of *g* means strong density regulation and the relative fitness-based model above is restored with *g* → ∞. Different strengths of regulation, *g_f_* and *g_r_*, are applied to the field and the refuge.

How a population evolves under this absolute fitness-based model was explored by the recursion of Eq. (3). Constant carrying capacity for the field (*K_f_* = 1000) and immediate redistribution over subpopulations (*m_fr_* = *m_rf_* = 0.5) were initially assumed. Four values of *g_f_* that represent the increasing strength of density regulation in the field were used (0.02, 0.2, 2, and 20). For the refuge, *g_r_* = 0 is assumed (as the model of seed bank is primarily considered), which makes it meaningless to specify *K_r_*.

The dynamics of a GMF allele was first explored. The fitness of *A*_1_ and *A*_2_ was given by *W*_1*f*_ = *e*^−0.5*s_t_*^, *W*_2*f*_ = *e*^0.5*s_t_*^, and *W_r_* = 1, where *st* is given by Eq. (1). Namely, the oscillation in absolute fitness of the two alleles is symmetric in the field. Under this fitness scheme *n*_1_, which initially oscillated around *2K_f_* (= 2,000), decreased towards an oscillation around *K_f_* (Fig. 2A). On the other hand, *n*_2_ increased from a small to large number that oscillates around *K_f_*. This approach of both alleles toward the relative frequency of 0.5 was faster with increasing *g*. (In Fig. 2A, the ranges of oscillation in *n*_1_ and *n*_2_ may appear to change little over time with *g* = 0.02 but they do converge close to 0.5 if iterations are done for a sufficiently large number of generations.) This observation is consistent with the storage effect arising in the relative fitness-based model, which is equivalent to this absolute fitness-based model with *g* → ∞.

**Figure 2.**
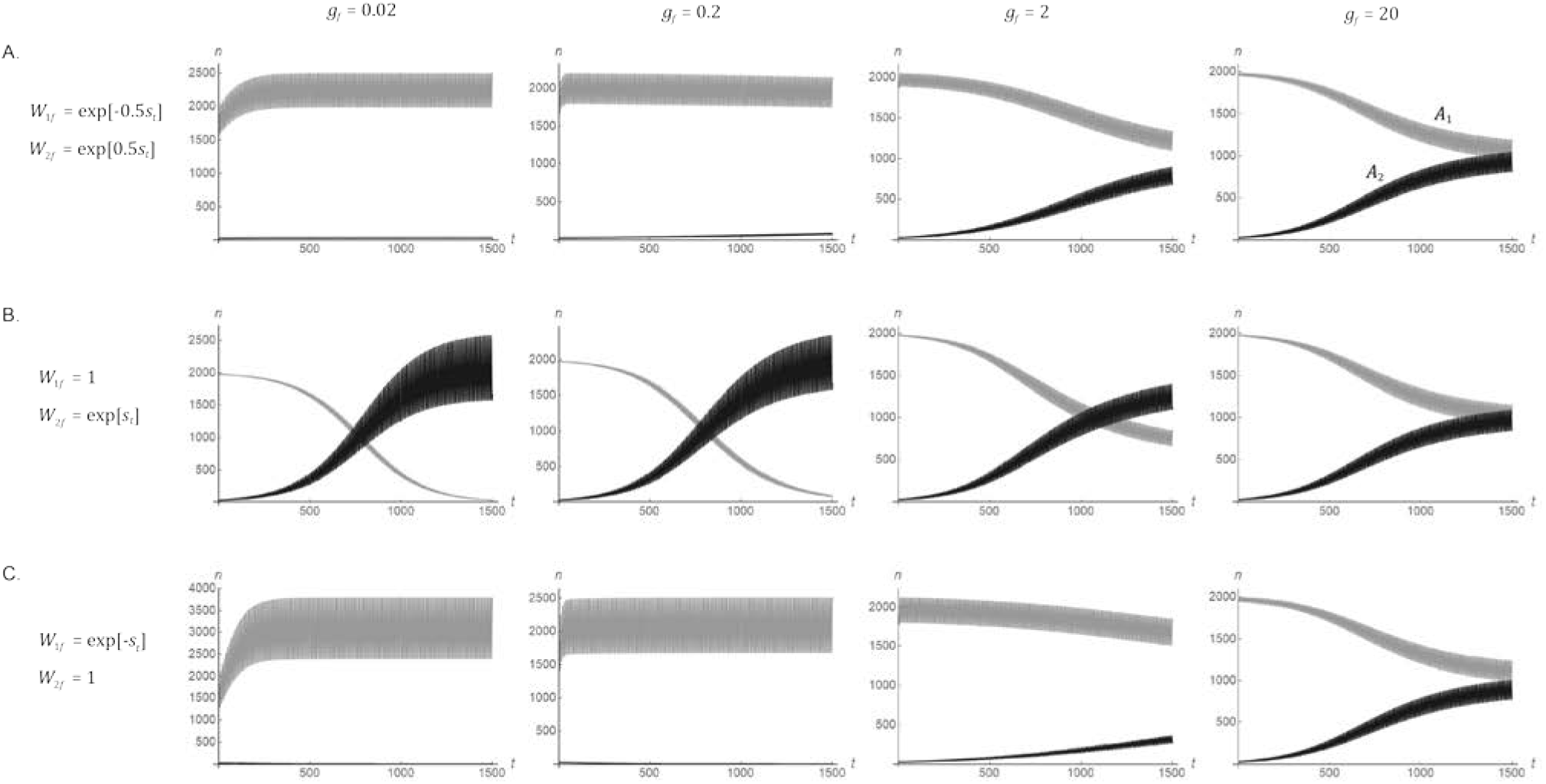
Changes in the absolute frequencies of *A*_1_ (*n*_1_ = *n*_1*f*_ + *n*_1*r*_; gray curves) and *A*_2_ alleles (*n*_2_ = *n*_2*f*_ + *n*_2*r*_; dark curves) in a population subdivided into the field and the refuge (*m_fr_* = *m_rf_* = 0.5), according to deterministic recursion of Eq. (3) starting from *n*_1*f*_ = *n*_1*r*_ = 990 and *n*_2*f*_ = *n*_2*r*_ = 10. Note that, as these graphs are unable to resolve curves oscillating very rapidly on the scale of time (in generations) shown above, they appear in the shape of bands. The field maintains a constant carrying capacity (*K_f_* = 1,000) with varying strengths (*g_f_*) of density regulation. Other parameters: *λ* = 10, *W_r_* = 1, *g_r_* = 0, *s_max_* = 0.3.

Next, the fitness scheme is changed to *W*_1*f*_ = 1 and *W*_2*f*_ = *e^s_t_^*, which keeps the relative fitness identical to the above scenario but lets the absolute fitness of *A*_2_ fluctuate more than that of *A*_1_. This dramatically changed the course of evolution (Fig. 2B). With strong density regulation (*g* = 20), the relative frequency of *A*_2_ approached 0.5, as before. However, under weaker density regulation, the absolute count of *A*_2_ continued to increase toward *2K_f_*, while that of *A*_1_ decreased toward zero. Similar results are obtained when the absolute fitness of both alleles fluctuate but the fluctuation of *A*_2_ is larger (e.g., *W*_1*f*_ = *e*^0.5*s_t_*^ and *W*_2*f*_ = *e*^1.5*s_t_*^). Namely, an allele increases in frequency by directional, not negative frequency-dependent, selection if its absolute fitness fluctuates more than that of the other allele. Switching the fitness oscillation of *A*_1_ and *A*_2_ (*W*_1*f*_ = *e*^−*s_t_*^ and *W*_2*f*_ = 1) confirmed this conclusion (Fig. 2C). Consequently, unless density regulation is strong, the population evolves toward experiencing a greater fluctuation in size.

This result – directional selection on an allele with larger fluctuation in absolute fitness – is again the consequence of assuming a GMF allele and the presence of refuge. Briefly, in the field, the arithmetic mean of *W*_2*f*_ = *e*^*s_t_*^ becomes increasingly larger than 1 as the oscillation of *s_t_* becomes larger in magnitude, while its geometric mean remains at 1. However, refuge reduces variance in the fitness of *A*_2_ in the total population, elevating its geometric mean (absolute) fitness above 1.

According to a simple mathematical analysis of this model at the limit of no density regulation (*g_f_* = 0), given in Appendix A, if the proportion of refuge is *r* in the total population and haploids are redistributed between the field and the refuge each generation (*m_fr_* = *r* and *m_rf_* = 1 − r), the increase in the number of an allele whose absolute fitness in the field fluctuates according to *W_f_* = *e^s_t_^* is given by

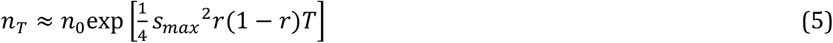

where *n*_0_ is the initial number and *T* is a multiple of *λ*. This approximates well the initial increase of *A*_2_ under weak density regulation given by exact recursion (Fig. S1). Eq. (5) shows that the geometric mean fitness of a GMF allele is maximized when the size of refuge is equal to that of the field (*r* = 0.5), and that, as the rate of increase is in the order of *s_max_*^2^, a significant change is expected only if the amplitude of fitness fluctuation is large (Bertram and Masel 2019). It is also shown in Appendix A that this increase does not occur for mutants with AMF.

**Figure S1.**
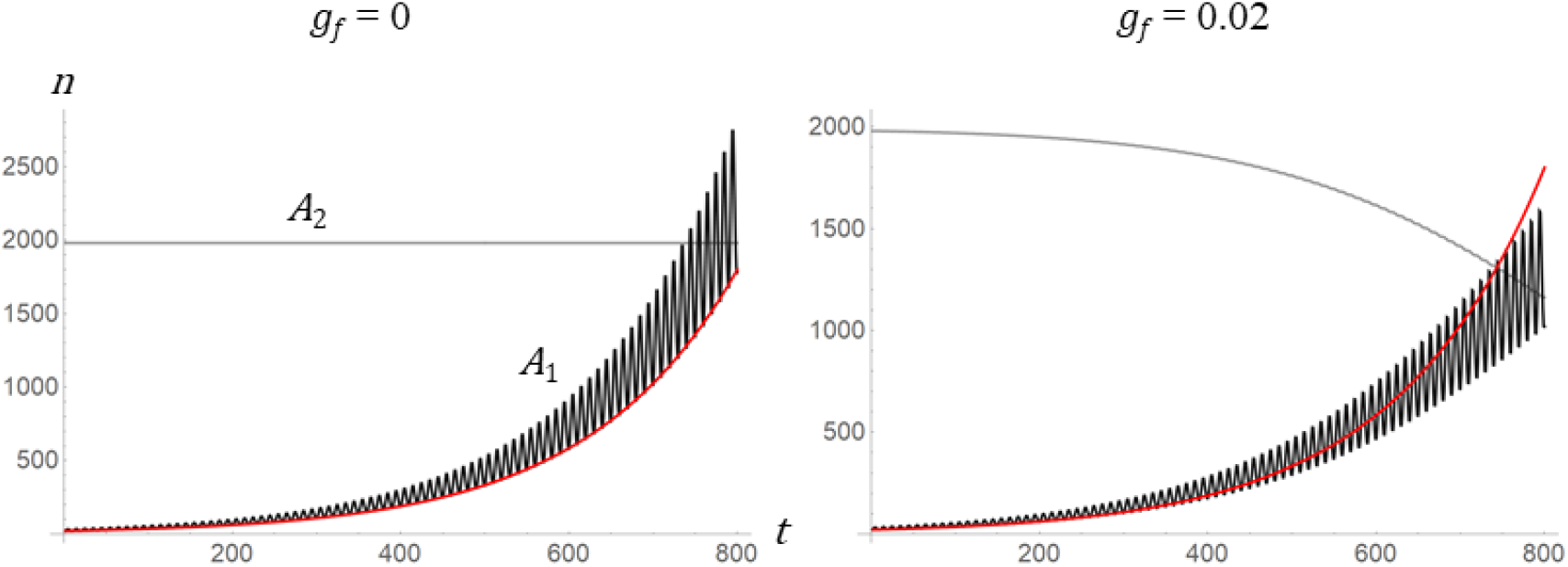
Comparison between the changes of *n*_1_ according to exact recursion (Eq. [3]; dark curves) and approximation by 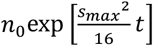 (red curves) in populations with *m_fr_* = *m_rf_* = 0.5 under weak density regulation (*g_f_* = 0 or 0.02). Fitness is given by GMF (*W*_1*f*_ = exp(*s_t_*) and *W*_2*f*_ = 1), with *s_max_* = 0.3. The changes of *n*_2_ by Eq. (3) are shown in gray curves.

The results shown in Fig. 2 can now be understood in terms of the “intrinsic growth rate” of a GMF allele under no density regulation described by Eq. (5). With positive (albeit weak) density regulation (*g_f_* > 0), the number of *A*_2_ cannot increase indefinitely. However, *A*_2_ outcompetes *A*_1_, whose absolute fitness fluctuates less (*W*_1*f*_ = 1 in Fig. 2B). Namely, the two alleles change in number according to their intrinsic growth rates before density regulation is applied, and their ratio determines the change in their relative frequencies. Under strong density regulation (*g_f_* ≫ 1), the relative frequency of *A*_2_ approaches a stable fluctuation around 0.5, at which *n*_1_ and *n*_2_ must fluctuate symmetrically so that their intrinsic rates of growth suggested by Eq. (5) become equal.

## Cyclic fluctuation of absolute fitness: oscillating carrying capacity

Next, to investigate whether the pre-existing fluctuation in population size affects the evolutionary dynamics of an allele with fluctuating fitness, the carrying capacity of the field was set to change cyclically, again with period 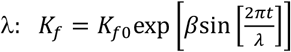, assuming *β* > 0 without the loss of generality. This models many species’ population cycles, which are commonly found to be the symmetric oscillations of density in the logarithmic scale (Myers 1988). Again, a GMF allele *A*_2_ whose fitness fluctuation is in or out of phase of the population size fluctuation arises at *t* = 0. The fitness scheme of *W*_1*f*_ = *e*^−0.5*s_t_*^ and *W*_2*f*_ = *e*^0.5*s_t_*^, for example, means that the *A*_2_ (*A*_1_) allele acts to amplify (dampen) the oscillation of population size beyond the level specified by the ecological factor *β* under weak density regulation. The recursion of Eq. (3) under this parameter set revealed that the *A*_2_ allele is positively selected to reach fixation, surprisingly, even under strong density regulation (*g_f_* = 20) (Fig. 3A). Similar rates of increase in *n*_2_ over *n*_1_ with *g_f_* > 1 were observed with other fitness schemes: (*W*_1*f*_, *W*_2*f*_) = (1,*e*^*s_t_*^) or (*e*^−*s_t_*^, 1) (Fig. 3B and 3C). On the other hand, if *A*_2_’s fitness fluctuation is out of phase with population size fluctuation (*W*_1*f*_ = 1 and *W*_2*f*_ = *e*^−*s_t_*^; Fig. 3D) positive selection for this allele under strong density regulation did not happen. Therefore, under strong density regulation, an allele is positively selected if its fluctuation in relative fitness is in phase with population size (carrying capacity) fluctuation. Further exploration revealed that the fixation of *A*_2_ allele however requires strong the fluctuation of *K_f_* for a given magnitude of fitness fluctuation, namely *β* > *s_max_* (Fig. S2). With relatively weaker fluctuation of population size, the long-term maintenance of polymorphism, the storage effect, was observed.

**Figure 3.**
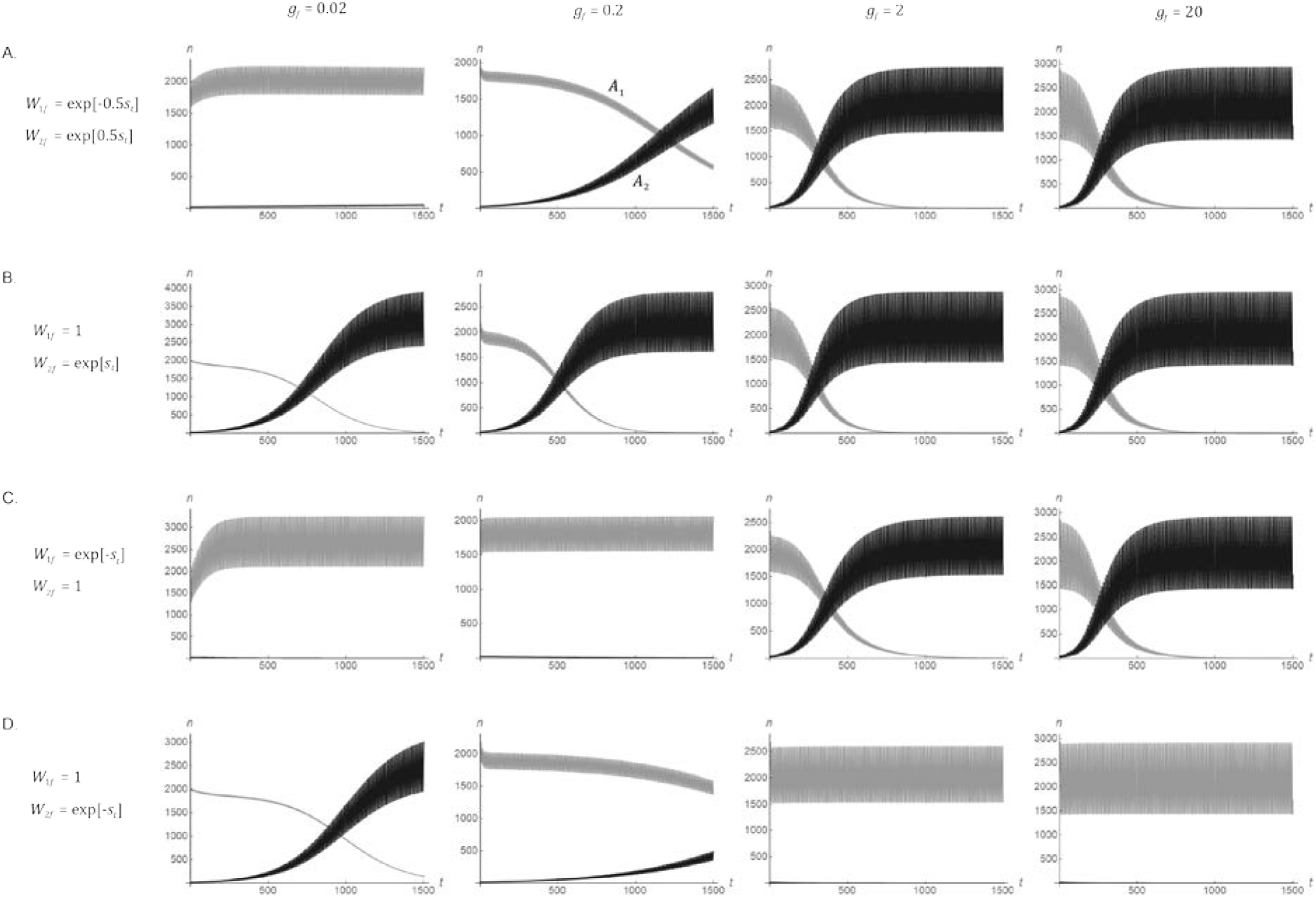
Changes in the absolute frequencies (*n*_1_ and *n*_2_) of *A*_1_ (gray curves) and *A*_2_ (dark curves) alleles under the cyclic fluctuation of carrying capacity in the field (*K*_*f*0_ = 1,000 and *β* = 0.5). Four fitness schemes that are classified as GMF [(*W*_1*f*_, *W*_2*f*_) = (*e*^−0.5*s_t_*^, *e*^0.5*s_t_*^) (A), (1, *e*^*s_t_*^) (B), (*e*^−*s_t_*^,1) (C), and (1, *e*^−*s_t_*^) (D)] were used. Other parameters are identical to those in Fig. 2.

With weak density regulation (*g_f_* = 0.02 and 0.2), the allele with a larger amplitude in absolute fitness fluctuation outcompetes the other, as in the case of constant carrying capacity (Fig. 2B and 3B), probably because the effect of (oscillating) carrying capacity on evolutionary dynamics diminishes as density regulation weakens. It should be noted again that, unless density regulation is very strong, the fixation of *A*_2_ leads to the increased magnitude of population size fluctuation compared to that when the population was fixed for *A*_1_ (Fig. 3).

**Figure S2.**
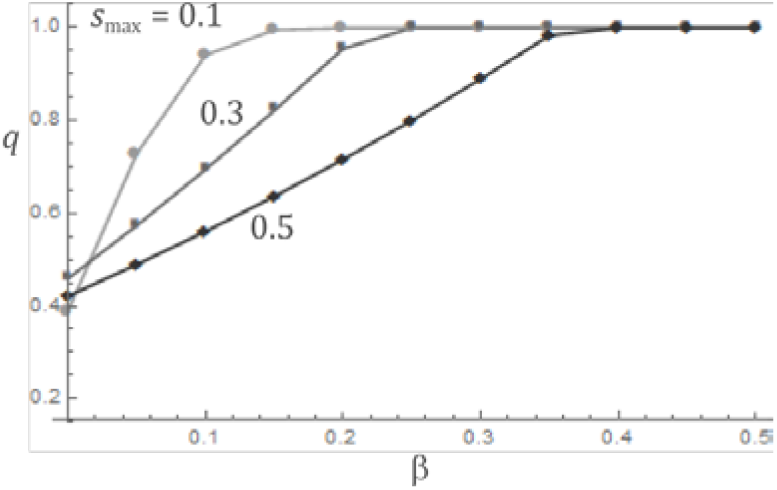
The relative frequency, *q*, of *A*_2_ allele (*W*_1*f*_ = 1, *W*_2*f*_ = *e*^*s_t_*^) at the 10,000^th^ generation of iterating eq. (3) with *g_f_* = 20. For *s*_max_ = 0.1, 0.3, and 0.5, increasing values of β, from 0 to 0.5 with increment of 0.05 were used. Other parameter values are identical to those in Fig. 3.

Assuming an AMF allele also resulted in diverse evolutionary outcomes with oscillating carrying capacity. As confirmed earlier, *A*_2_ failed to increase in frequency when *W*_1*f*_ = 1 and *W*_2*f*_ = 1 + *s_t_* with a constant carrying capacity of the field (Fig. 4A), as its geometric mean fitness in the field alone is less than 1. However, under cyclic oscillation of *K_f_* (β = 0.5) and strong density regulation (*g_f_* = 2 or 20), the frequency of *A*_2_ increased by positive selection. With *s*_max_ = 0.3, *A*_2_ did not reach fixation but remained polymorphic with *A*_1_ in the long run (Fig. 4B), suggesting that a form of balancing selection arose. With smaller *s*_max_ (= 0.2), however, *A*_2_ finally reached fixation (Fig. 4C). Again, these results depend on the condition that the fitness oscillation of *A*_2_ is in phase with population size oscillation in the field. If fitness oscillation was out of phase (*W*_1*f*_ = 1 and *W*_2*f*_ = 1 − *s_t_*), positive selection on *A*_2_ did not occur (Fig. 4D).

**Figure 4.**
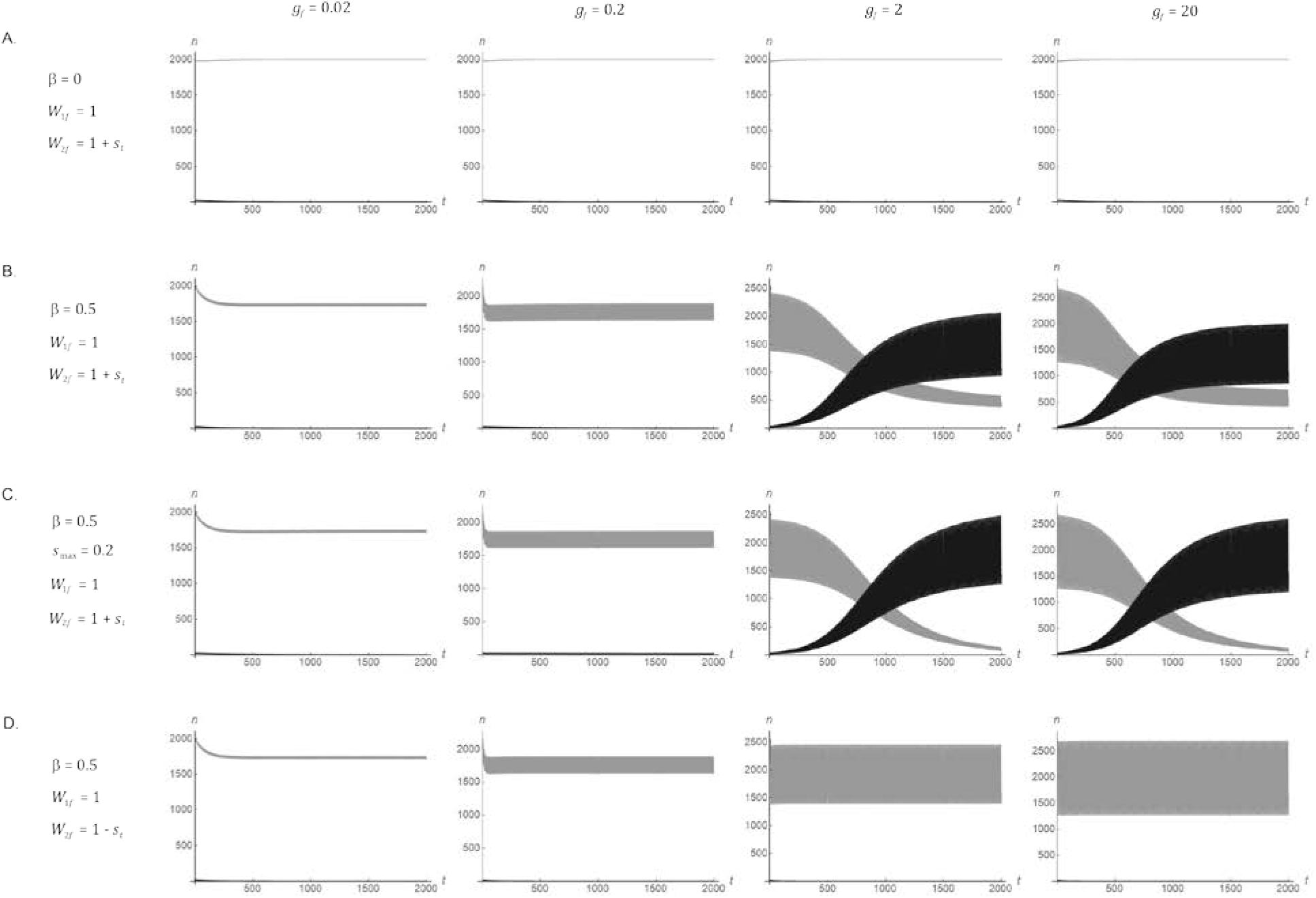
Changes in the absolute frequencies (*n*_1_ and *n*_2_) of *A*_1_ (gray curves) and *A*_2_ (dark curves) alleles when the fitness scheme of *A*_2_ is classified as AMF: (*W*_1*f*_, *W*_2_*f*) = (1,1 + *s_ŧ_*) (A, B, C), and (1,1 − *s_t_*) (D). Carrying capacity in the field is either constant (β = 0; A) or oscillating (β = 0.5; B, C, D). The magnitude of fitness fluctuation, *s*_max_, is either 0.3 (A, B, D) or 0.2 (C). Other parameters are identical to those in Fig. 3. When much longer iteration (> 20,000) was done for the parameter set shown in panel B with *g_f_* = 20, *n*_1_ and *n*_2_ were observed to be oscillating at values that have changed little from those observed at generation 2,000.

## Two-season model for fitness and population size fluctuation

For the general understanding of positive selection generated under the oscillation of carrying capacity, revealed above by numerical iterations of eq. (3) for a limited set of parameters, a simpler model at the limit of strict density regulation is analyzed. A cyclic environment with λ = 2 is assumed, in which each period is made up of a “good” season (the first generation) and a “bad” season (the second generation). The carrying capacity of the field then alternates between *K*_G_ and *K*_B_, where *K*_G_ ≥ *K*_B_. The absolute fitness of *A*_1_ and *A*_2_ individuals in the field are assumed to be *W*_1G_ and *W*_2G_ in a good season and *W*_1B_ and *W*_2B_ in a bad season. In the refuge, *W_r_* = 1. Strict density regulation in the field but no density regulation in the refuge (*g_f_* → ∞ and *g_r_* = 0) is assumed. Then, assuming *m_fr_* = *r* and *m_rf_* = 1 − *r*, this population reaches a demographic equilibrium in which the total population size alternates between *N*_G_ (at the end of a good season) and *N*_B_ (at the end of a bad season). In Appendix B, it is shown that the number of haploid individuals carrying *A*_2_ allele changes from *n*_2_ to *n*″_2_ in one cycle, where

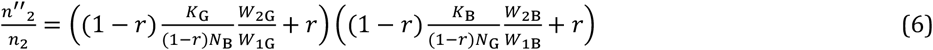

approximately, if *n*_2_ is far smaller than *n*_1_. Eq. (6) shows that, as expected under strict density regulation, the evolutionary change of *n*_2_ is dependent on the relative, not absolute, fitness of alleles. However, because the absolute, not relative, frequency of *n*_2_ is tracked in the derivation, the growth rates of population size in the field should be combined with the relative fitness to determine its evolutionary change; the field grows from (1 − *r*)*N*_B_ to *K*_G_ during the good season and shrinks from (1 − *r*)*N*_G_ to *K*_B_ during the bad season.

Now, *A*_2_ is assumed to be a GMF allele. The relative fitness between two alleles is reversed each season: 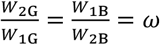. Then, the change in the number of *A*_2_ when it is rare is given by

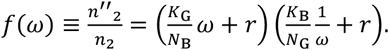

It can be easily shown that *f*(1) = 1, as expected for a neutral allele. Solving *f* (*ω*) = 1 and analyzing the derivative of *f* (*ω*) shows that

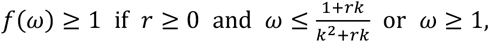

where 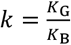, and *f*(*ω*) < 1 otherwise (Appendix B). Therefore, positive selection on *A*_2_ allele requires the presence of refuge. If *K*_G_ = *K*_B_ (no change in population size), *f*(*ω*) > 1 for all *ω* ≠ 1. Since *f*(ω^−1^) describes the change of *A*_1_ frequency when *A*_1_ is rare, this result means that both *A*_1_ and *A*_2_ alleles are positively selected when they are rare. This symmetry creates negative frequency-dependent selection—the storage effect. However, if *K_G_* > *K_B_* and 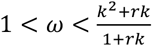, *A*_2_ is positively selected but *A*_1_ is not, even if *A*_1_ is rare. This asymmetry means that *A*_2_, whose fitness fluctuates in the same direction as (i.e., in phase with) population size fluctuation, is under directional selection and will eventually reach fixation. For example, if *K*_G_ = 2*K*_B_, an allele that is between 1 and 2.5 times advantageous over the other allele during a good season but inversely disadvantageous during a bad season is positively selected until it reaches fixation. Lastly, if 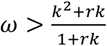, both *A*_1_ and *A*_2_ alleles are positively selected when they are rare, thus yielding the storage effect again: fitness fluctuation is much stronger than population size fluctuation so that the qualitative behavior of system is identical to that with no population size fluctuation.

In case of an AMF allele, we may assume 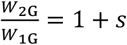 and 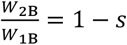. The change in the number of *A*_2_ when it is rare is given by

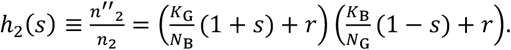

Solving *h*_2_(*s*) ≥ 1 gives 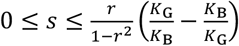. Therefore, unless *s* is too large relative to the magnitude of population size fluctuation (*K*_G_/*K*_B_), *A*_2_ allele whose fitness fluctuation is in phase with population size fluctuation invades the population. At the same time, the condition for the *A*_1_ allele invading a population fixed with *A*_2_ is given by

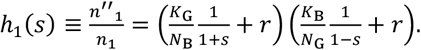

For 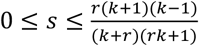, where 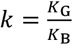, *h*_1_(*s*) ≤ 1. Note that 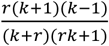 is an increasing function of *k* (> 1). Therefore, with *s* that is small for a given magnitude of population size fluctuation, the condition for *A*_2_ increasing in frequency to fixation is satisfied. However, with 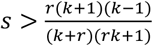, *h*_1_(*s*) > 1: *A*_1_ can invade the population. Therefore, with an intermediate strength of fluctuating selection 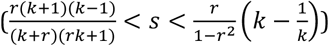, polymorphism with *A*_1_ and *A*_2_ should be maintained. Parameter ranges leading to different evolutionary outcomes under the assumption of AMF are graphically summarized in Fig. 5.

**Figure 5.**
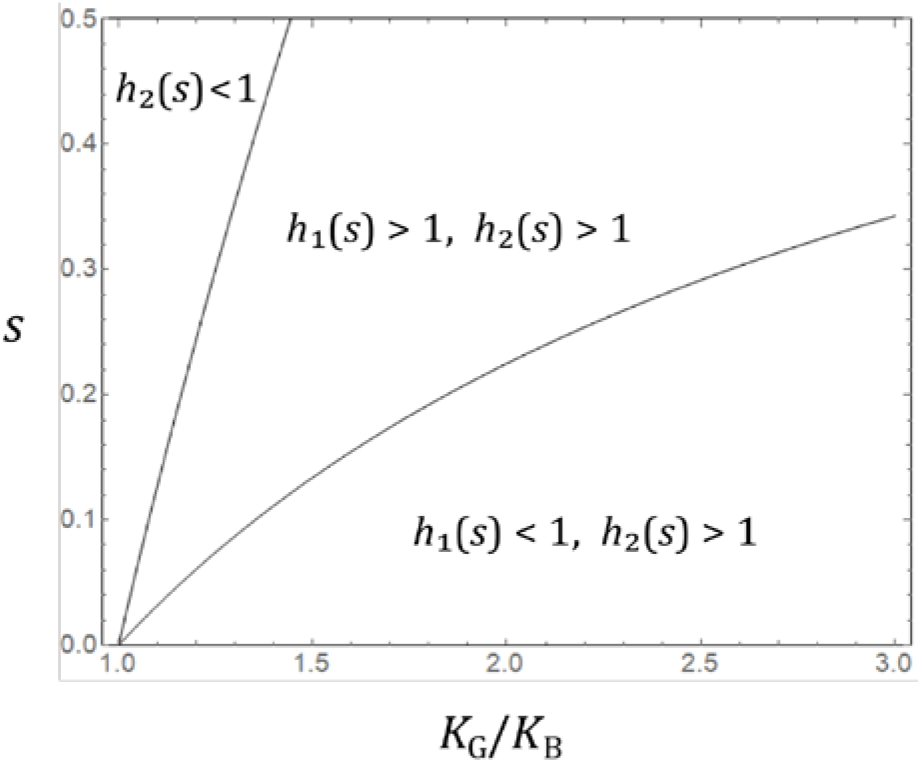
Domains of *s* and *K*_G_/*K*_B_ in which negative selection (*h*_2_(*s*) <1), balancing selection (*h*_1_(*s*) >1, *h*_2_(*s*) >1), and directional selection (*h*_1_(*s*) <1, *h*_2_(*s*) >1) on the derived allele *A*_2_ occurs in the two-season model (*r* = 0.5) of AMF and population size fluctuation.

## Random fluctuation of absolute fitness

Finally, it was examined whether evolutionary dynamics produced in the absolute fitness-based model also arise when the absolute fitness fluctuates randomly, rather than cyclically. Iteration of Eq. (3) was performed using the same set of parameters above. However, at each generation a random integer *t*\* was drawn from 1 to *λ*. Then, the carrying capacity and selection coefficient were given by *K_f_* = *K*_*f*0_exp[*β*sin[2*πt*\*/*Λ*]] and *s_t_* = *s*_max_sin[2*πt*\*/*Λ*]. Therefore, if *β* > 0 and the fitness of the *A*_2_ allele is *W*_2*f*_ = *e*^*s*^_*t*_ or 1 + *s_t_*, *K_f_* and *W*_2*f*_ fluctuate in the same direction; it models an allele that, for example, leads to the immediate consumption of all available resources, the amount of which changes randomly, rather than even allocation of resources over successive generations. For each parameter set, 100 iterations were made and the mean frequency trajectory was obtained (Fig. S2).

Random fitness fluctuation yielded all major results observed with cyclic fitness fluctuation: GMF alleles positively selected and then maintained at intermediate frequency under strong density regulation with constant carrying capacity (Fig S2A, B), GMF alleles with wider fluctuation in absolute fitness positively selected toward fixation under weak density regulation with or without population size fluctuation (Fig. S2B, C), GMF and AMF alleles whose fitness fluctuates in the same direction as population size positively selected toward fixation under strong density regulation (Fig. S2C, D), and, with a smaller magnitude of population size fluctuation (β = 0.3 instead of 0.5), AMF alleles positively selected and then maintained polymorphic under strong density regulation (Fig. S2E). The polymorphic equilibria reached in the last cases were confirmed in longer iterations (Fig. S3).

**Figure S2.**
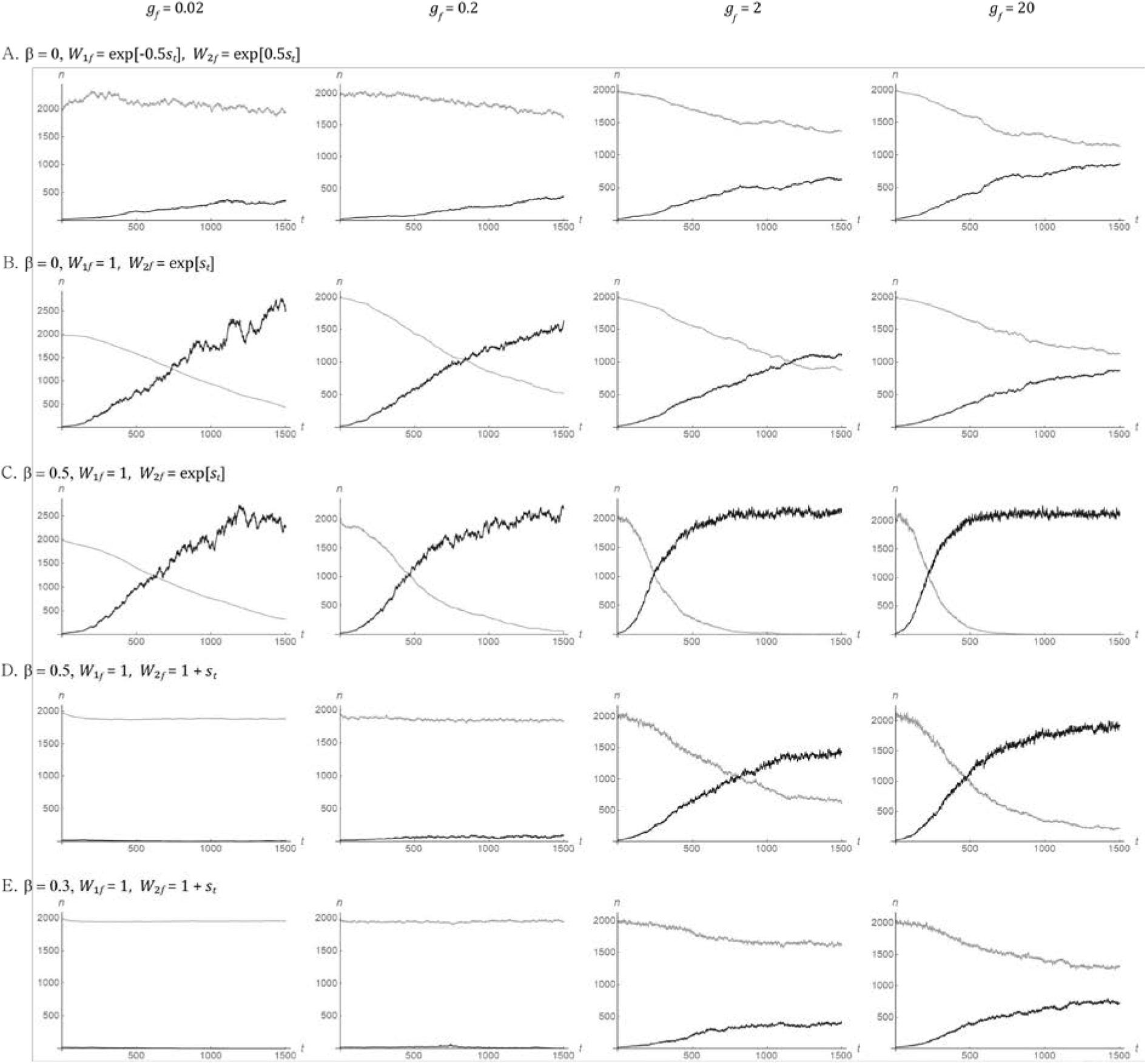
Changes in the absolute frequencies (*n*_1_ and *n*_2_) of *A*_1_ (gray curves) and *A*_2_ (dark curves) alleles under the random fluctuation of carrying capacity on the field (*K*_*f*0_ = 1,000, *K_f_* = *K*_*f*0_exp[*β*sin[2*πt*\*/*Λ*]], and *s_t_* = *s_max_*sin[2*πt*\*/*Λ*], where *t*\* is a random integer between 1 and *λ* drawn each generation). Other parameters are identical to those in Fig. 3.

**Figure S3.**
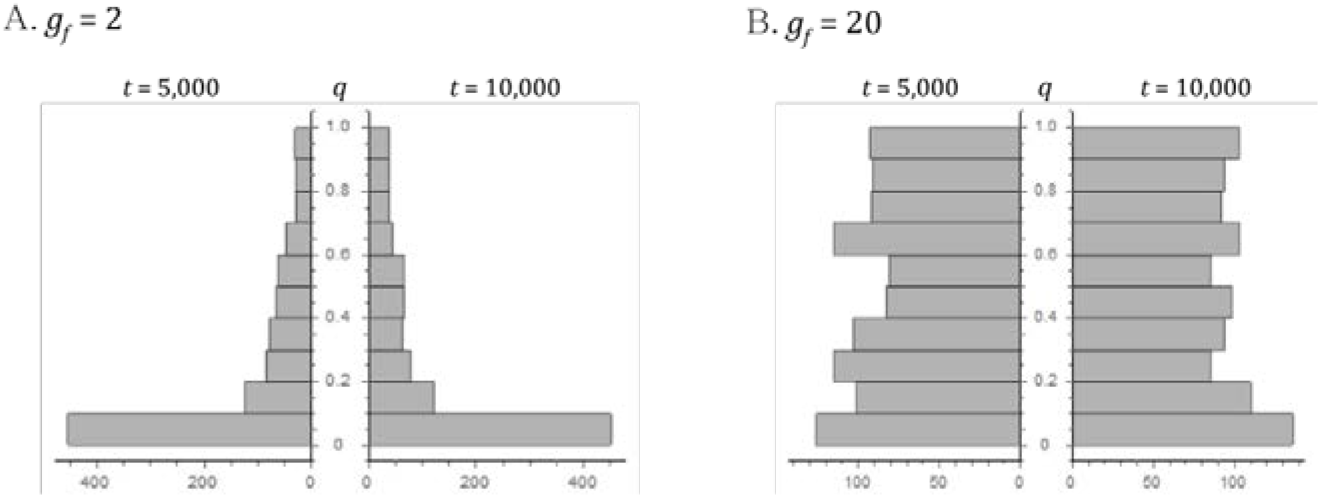
Paired histograms showing the distribution of *A*_2_ frequency (*q*) at the 5,000^th^ and 10,000^th^ generation in the iteration of eq. (3) with random fitness fluctuation, corresponding to *g*_f_ = 2 (A) and 20 (B) of Fig. S2E. 1,000 iterations in total were performed for each parameter set.

## DISCUSSION

The model of fluctuating selection in the presence of refuge was modified in terms of absolute fitness to accommodate the joint dynamics of population size and allele frequency changes. The general importance of investigating the coupled demographic and evolutionary dynamics was already well demonstrated. The outcome of natural selection under a changing population size can differ from that under a constant population size, as shown in the case of the fixation probability of beneficial alleles (Otto and Whitlock 1997) or the nature of fitness that is maximized (Sæther and Engen 2015). Conversely, various demographic changes can occur as the result of natural selection, as in the case of evolutionary rescue or trapping (Carlson et al. 2014). Furthermore, as shown in studies using simple mathematical models (Pimentel 1961; Charlesworth 1971; Roughgarden 1971; Lion 2018), demographic and evolutionary dynamics occur in a tightly-coupled process, which is described as an eco-evolutionary feedback (Kokko and Lopez-Sepulcre 2007; Govaert *etal*. 2019).

This study found that the model of fluctuating selection with refuge makes drastic changes in prediction if it is built upon absolute instead of relative fitness. First, under weak density regulation, a GMF allele that causes a wider fluctuation in population size is under positive directional selection instead of negative frequency-dependent selection. The latter, referred to as the storage effect, has long been proposed as a mechanism for promoting species or genetic diversity (Chesson and Warner 1981; Bertram and Masel 2019). It was also clarified that AMF alleles are not subject to either balancing or directional selection in the presence of refuge if the carrying capacity of the population (the field) is constant over time. These results therefore calls for significant revisions in conclusions drawn from previous studies on the storage effect.

Second, more surprisingly, preexisting fluctuation in population size was found to select an allele, regardless of whether it is GMF or AMF, if its relative fitness fluctuates in the same direction with population size under strong density regulation. In case of derived AMF alleles, either negative, balancing, or directional selection on them occurs depending on the magnitude of fitness fluctuation relative to population size fluctuation. To my knowledge this type of positive selection, acting on either quasi-neutral (GMF) or deleterious (AMF) alleles outside the refuge, induced by demographic changes was not discovered before. The emergence of balancing selection – leading to the long-term oscillation of an AMF allele at intermediate frequencies – driven by population size fluctuation is particularly remarkable. Although this mechanism also requires the presence of a refuge, it should be distinguished from the storage effect because the latter arises as a larger arithmetic mean of a derived allele’s relative fitness is converted into a larger geometric mean due to the variance-reducing effect of refuge. Here, the derived allele is not conferred a larger arithmetic mean. Mutations to AMF alleles may arise frequently as many life history traits are subject to trade-off given a finite resource. Then, as the whole genome is affected by population size fluctuation, positive selection on AMF alleles can arise at many loci independently as long as their fitness fluctuation is in phase with demographic fluctuation. This is a potential explanation for the seasonal oscillation of allele frequencies at more than 1,000 loci in North American *Drosophila melanogaster* populations (Bergland *et al*. 2014). While further refinement of theory is likely needed and the ecology of these fruit flies needs to be elucidated before confirming whether this novel mechanism of balancing selection can operate in the actual population, the major requirements for the theory – clear seasonality in the temperate region and the pupal stage that can serve as a refuge – appear to be satisfied in these populations.

The fluctuation of population size in response to either a randomly or cyclically fluctuating environment is observed in many plant and animal species, particularly those reproducing in multiple generations over one seasonal cycle. Classical results in population genetics predict that a genotype causing less fluctuation in fitness is favored under such an environment (Gillespie 1977; Lande et al. 2009). Namely, studies suggested that, given a mutation increasing fitness in one phase of a temporally changing environment and decreasing it in another phase as a trade-off, it is positively selected if the trade-off occurs in a direction that decreases the amplitude of population size fluctuation. Such a mutation, for example, would change the allocation of resources toward a season unfavorable for the survival and/ or reproduction of a species. This study, however, showed that natural selection can occur in the opposite direction if a population is already partially protected from selection. Under weak or moderate density regulation, even if the population size is initially constant over time, a GMF allele causing population size fluctuation is positively selected toward fixation (Fig. 2B). If the population size is already fluctuating due to external forces (a variable environment) and/or due to the fixation of above-mentioned allele at another locus, an allele causing a wider fluctuation of absolute fitness can invade and therefore amplify the fluctuation in population size under moderate density regulation (cases with *g_f_* = 0.02 and 0.2 in Figs. 3 and 4). Such an allele, for example, allocates further resources to a period during which the rate of growth and reproduction is already high, which inevitably causes further deterioration of fitness during other periods. Assuming a large number of loci at which mutations affecting life history traits can arise, one can imagine that selection at multiple loci can reinforce each other until a very wide fluctuation in population size arises.

In the theory of evolutionary bet-hedging (Seger and Brockmann 1987; Philippi and Seger 1989), a refuge from selection such as a seed bank emerges in a reproductive strategy that may decrease the arithmetic mean but increase the geometric mean of fitness by reducing the variance of reproductive success. Namely, plants’ delayed germination or insects’ diapause can be the result of natural selection toward less severe fluctuations in fitness. In this study, however, refuge from selection is given as a fixed demographic parameter in the model under which selection acts on variants causing different patterns of fitness fluctuations. Given the current results, findings in the previous investigation on evolutionary bet-hedging may be interpreted differently. For example, by fitting the field observation of annual desert plants to their mathematical model, which closely resembles Eq. (3), Gremer and Venable (2014) showed that species experiencing wider fluctuation in fitness after germination have greater proportions of their population remaining in the seed bank. This supports the hypothesis that delayed germination evolved as a bet-hedging strategy. However, it could be other life history traits that evolved in those species; in species with a larger probability of staying in the refuge (larger seed bank), variants causing wider fluctuations in fitness in the field (after germination) could have been selected, as suggested in this study.

The storage effect discovered in the previous studies and evolution toward amplified fluctuation in population size discovered in this study depend on the occurrence of mutations that cause fitness fluctuations without decreasing their geometric mean relative to the ancestral allele. Because these GMF alleles are required to produce more offspring than their ancestral alleles within an environmental cycle, they may be considered as a type of beneficial allele. Such mutations probably arise less frequently than mutations generating AMF alleles, which can arise for example by simply changing resource allocation over different seasons. However, there can be many biological scenarios under which mutants’ fluctuating fitness arise in a way to preserve the geometric mean. For example, imagine that the body size of a haploid individual carrying *A*_2_ allele relative to *A*_1_ is 1 + *x* (> 1) in one season and 1 - *x* in the next season, compatible with the trade-off of finite resource. Assume further that, when they encounter a predator with probability *y* for both seasons, the predator may always choose the smaller individual to attack. Then, the relative fitness of *A*_2_ becomes 1/(1 − *y*) in the first season and 1 − *y* in the second season, making *A*_2_ the GMF allele.

In conclusion, a simple mathematical model clearly revealed novel eco-evolutionary dynamics, in which an evolutionary change leads to a demographic change (i.e. positive selection on a variant causing a wider fluctuation in population size) and a demographic change leads to an evolutionary change (i.e. population size fluctuation selecting a variant with in-phase fitness fluctuation). Such two-way interaction between ecology and evolution cannot be discovered if demography is given as a fixed parameter as in the conventional models of population genetics.

## Acknowledgement

This study was supported by the Research Foundation (NRF) grant 2020R1A2C1009261 funded by the Korean government.

## Appendix A. Growth rate under fluctuating absolute fitness in the presence of refuge

At the limit of no density regulation (*g*_f_ = 0) the number, *n*, of haploid individuals carrying a given allele changes independently of the other allele. Assuming that *m_fr_* = *r* and *m_rf_* = 1 − *r* and that this allele has the absolute fitness of *W_f_* and *W_r_* in the field and in the refuge, respectively, *n* changes to *n*((1 − *r*)*W_f_* + *rW_r_*) over a generation. Starting from *n_0_* the number of individuals after a single cycle of fitness fluctuation is given by

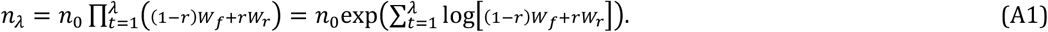

First, the absolute fitness of a GMF allele can be given by 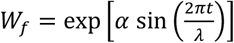, where *α* determines the amplitude of fluctuation, and *W_r_* = 1. Then,

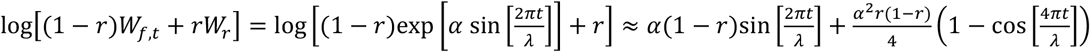

after Taylor expansion around *α* = 0 up to the second order. Plugging the above into eq. (A1) yields,

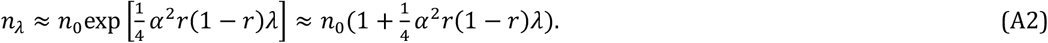

Namely, the absolute number of individuals increases as long as their absolute fitness fluctuates in the field, when no such change is expected in the field or refuge in isolation (⊓_t_ *W_f_* = 1 and *W_r_* = 1).

Second, in the case of an AMF allele, absolute fitness can be given by 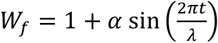 and *W_r_* = 1. Using Taylor approximation again,

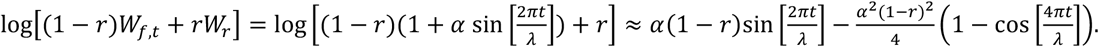

Then,

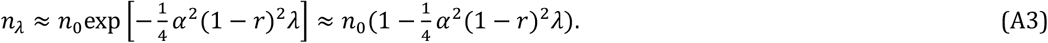

A refuge therefore cannot turn the negative growth rate of a subpopulation carrying an AMF allele from a negative to a positive value. A wider fluctuation of fitness accelerates this decrease.

## Appendix B. Demography-driven selection in the two-season model under strict density regulation

One environmental cycle consists of two generations, which correspond to two seasons, good and bad, with the carrying capacity of the field given by *K*_G_ and *K*_B_, respectively. At the start of each generation (season), haploid individuals migrate to the field and the refuge with probability 1 - *r* and *r* regardless of their current location. At the end of bad season there are *N*_B_ individuals. During the good season, the population size of the field changes from (1 − *r*)*N*_B_ to *K*_G_ while that of the refuge remains at *rN*_B_ since *W_r_* = 1 is assumed. Total population size at the end of a good season is therefore *N*_G_ = *K*_G_ + *rN*_B_. Similarly, *N*_B_ = *K*_B_ + *rN*_G_. Therefore, *N*_G_ = (*K*_G_ + *rK*_B_)/(1 − *r*^2^) and *N*_B_ = (*K*_G_ + *rK*_B_)/(1 − *r*^2^). Let *n*_1_ and *n*_2_ be the number of *A*_1_ and *A*_2_ individuals at the end of a bad season (therefore, *n*_1_ + *n*_2_ = *N*_B_). Then, *n*_2_ increases to

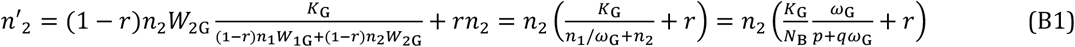

at the end of good season, where *ω*_G_ = *W*_2G_/*W*_1G_ is the relative fitness of *A*_2_ in the field in good season and *q* = 1 − *p* = *n*_2_/(*n*_1_ + *n*_2_). Using also

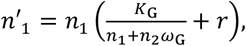

the number of *A*_2_ individuals at the end of the following winter generation changes to

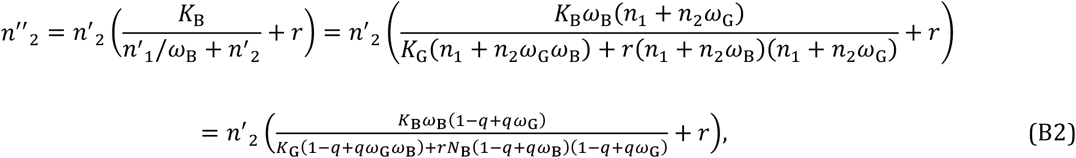

where *ω*_B_ = *W*_2B_/*W*_1B_. When *A*_2_ is rare (*q* → 0) the change in its absolute frequency after one cycle is obtained from eqs. (B1) and (B2):

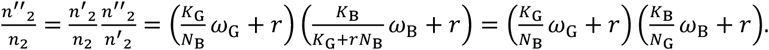

If *ω*_G_ = *ω*_B_ = 1, the above becomes

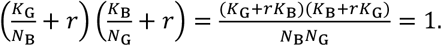

Then, if *ω*_G_ = 1/*ω*_B_ = *ω*, the condition for *A*_2_ allele invading the population 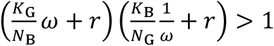 is rewritten as

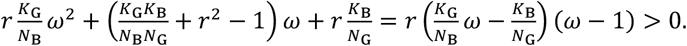

If *ω*_G_ = 1 + *s* and *ω*_B_ = 1 − *s*, the condition for the invasion of *A*_2_ is

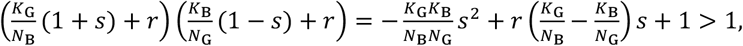

which is satisfied for

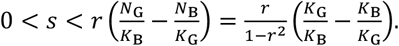

At the same time, the condition for the invasion of *A*_1_, 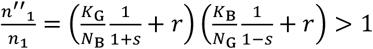 is rearranged to

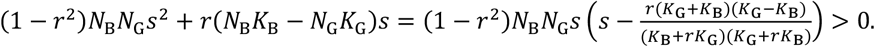

